# A two-step probing method to compare lysine accessibility across macromolecular complex conformations

**DOI:** 10.1101/448605

**Authors:** Andrew J. MacRae, Patricia Coltri, Eva Hrabeta-Robinson, Robert J. Chalkley, A.L. Burlingame, Melissa S. Jurica

## Abstract

Structural models of multi-megadalton molecular complexes are appearing in increasing numbers, in large part because of technical advances in cryo-electron microscopy realized over the last decade. However, the inherent complexity of large biological assemblies comprising dozens of components often limits the resolution of structural models. Furthermore, multiple functional configurations of a complex can leave a puzzle as to how one intermediate moves to the next stage. Orthogonal biochemical information is crucial to understanding the molecular interactions that drive those rearrangements. We present a two-step method for chemical probing detected by tandem mass-spectrometry to globally assess the reactivity of lysine residues within purified macromolecular complexes. Because lysine side chains often balance the negative charge of RNA in ribonucleoprotein complexes, the method is especially powerful for detecting changes in protein-RNA interactions. Probing the *E. coli* 30S ribosome subunit showed that the reactivity pattern of lysine residues quantitatively reflects structure models from X-ray crystallography. We assessed differences in two conformations of purified human spliceosomes. Our results demonstrate that this method supplies powerful biochemical information that aids in functional interpretation of atomic models of macromolecular complexes at the intermediate resolution often provided by cryo-electron microscopy.

## INTRODUCTION

Cellular macromolecular complexes, including ribonucleoproteins (RNPs) like the ribosome, the spliceosome and telomerase, often contain many moving parts, making it challenging to establish the mechanisms that control their molecular motions. Recent cryoelectron microscopy (cryo-EM) technological advances are now yielding the first models of complicated macromolecular assemblies previously refractory to structure analysis. Many of these assemblies, such as the spliceosome, telomerase and transcription initiation complexes (1-21), have also been captured in different functional conformations. While such structures provide important clues to the molecular mechanisms of these cellular machines, the intermediate resolution ranges (3-10 Å) of the cryo-EM maps cannot predict interaction details.Furthermore, cryo-EM models, like X-ray crystallographic structures, are derived from stable conformations that often represent only one of many intermediate states within a multi-step process and/or assembly pathway. Additional means of measuring conformational changes of complexes remains essential to wholly describing how such complexes assemble and function.

Partnered with chemical modification, mass spectrometry (MS) has emerged as a critical tool for gleaning structural information from macromolecular complexes. Tandem MS/MS peptide sequencing is used to locate the position of chemical crosslinks to help define protein interactions (19). It can also be used to quantify hydrogen/deuterium exchange to estimate solvent accessibility and flexibility of protein regions (22). Similarly, chemical probing (covalent labeling) exploits differential reactivity to reagents that non-specifically modify protein side chains, such as hydroxy radicals, or that target specific amino acids depending on a residues exposure to solvent vs. participation in interactions within the macromolecule (23,24).

Several reagents are available for amino acid probing with well-defined chemistries (23), but for RNPs, the reactivity of lysine side chains to a probing reagent is particularly informative. Lysine reactivity to acetylation reagents reflects both solvent accessibility and availability of the lone pair electron of the uncharged ɛ-nitrogen as a nucleophile. When positively charged, the primary amine of lysine side chains often help counteract and position the negatively charged phosphate backbone of nucleic acids (25). Amino acid probing is commonly carried out with well-behaved individual proteins that can be purified in high (> nanomolar) concentrations, and with the goal of examining surface topology and protein/ligand interactions (23). Extending chemical probing methods to multiple conformations of larger and often dynamic assemblies, including RNPs that are often only available at low picomolar amounts, brings a unique set of challenges.

To address the challenges related to probing multiple conformations of the same RNP we developed a chemical probing strategy that includes two lysine acetylation steps. Peptides for MS/MS analysis are usually generated by trypsin digest. Identifying a change in modification state depends on analyzing the same peptide from different samples, which is problematic because acetylated lysines are not susceptible to trypsin digestion. Therefore, we added a second acetylation step under denaturing conditions, which links a deutero-acetyl group to lysines that in the intact / assembled RNP were protected in the first modification reaction. The second acetylation step ensures that the same tryptic peptides are generated across different samples.Furthermore, the presence of acetyl vs. deutero-acetyl modification differentiates lysine reactivity in the intact macromolecule.

Here we describe the two-step acetylation protocol and its application to two different RNP complexes: 30S ribosome subunit from *E. coli* and spliceosomes assembled in nuclear extract from human cells. Our results indicate that lysine reactivity is indeed reflective of complex structure, and can be used to characterize conformational changes in macromolecular complexes. Probing data can also serve as a guide for more mechanistic analysis. For example, we recently showed a functional role for specific lysine residues in the spliceosome protein Prp8, which we identified by their altered accessibility before and after assembly of they active site (26). Although we have focused on RNPs, the method is compatible with single proteins and protein complexes.

## MATERIAL AND METHODS

### 30S Ribosome Subunit Reconstitution

Purified 30S *E. coli* ribosomal RNA and ribosomal proteins (provided by the Noller lab at UCSC) in 10 mM Tris-HCl pH 7.5, 100 mM NH_4_Cl, 10 mM MgCl_2_, 6 mM BME were assembled into 30S subunits via incubation at 42°C for 15 mins. After reconstitution, 30S ribosome subunits were exchanged into 20 mM HEPES pH 7.5, 100 mM KCl, 10 mM MgCl_2_ and 1 mM DTT via Slide-A-Lyzer 10 KDa mini-dialysis device (Thermo Fisher Scientific) for 5 hours at 4°C.

### In vitro spliceosome assembly and purification

Catalytic and post-catalytic pre-mRNA substrates are derivatives of the AdML transcript, are tagged with three MS2 sites in either the intron or at the 3′ end, respectively. T7 runoff transcription was used to generate G(5′)ppp(5′)G-capped radiolabeled pre-mRNA, which was gel purified and preincubated with a 50-fold excess of MS2-MBP fusion protein. Catalytic spliceosomes arrested after first-step chemistry were assembled with a pre-mRNA containing a mutant GG 3′ splice site as previously described (27). Post-catalytic spliceosomes arrested after second-step chemistry were assembled with a pre-mRNA containing a truncated 3 exon as previously described (28).

Both complexes were assembled in *in vitro* splicing reactions containing 10 nM premRNA, 80 mM potassium glutamate, 2 mM magnesium acetate, 2 mM ATP, 5 mM creatine phosphate, 0.05 mg/mL tRNA, and 40% HeLa cell nuclear extract at 30°C for 60 minutes. The reactions were fractionated by size exclusion chromatography using Sephacryl S-400 resin (GE Healthcare) in 20 mM HEPES pH 7.9, 150 mM potassium chloride, 5 mM EDTA, 0.05% v/v NP-40, and then bound to amylose resin. After washing in the same buffer, spliceosomes were eluted from the amylose resin in 20 mM HEPES pH 7.9, 150 mM potassium chloride, 5 mM EDTA, and 1 μM maltose.

### Chemical probing by two-step acetylation

For the first acetylation step, one to five picomoles of reconstituted 30S *E. coli* ribosome subunits (20 mM HEPES pH 7.5, 100 mM KCl, 10 mM MgCl_2_ and 1 mM DTT), isolated 30S ribosome proteins (20 mM HEPES pH 7.5, 100 mM KCl, 10 mM MgCl_2_ and 1 mM DTT) or human spliceosomes (20 mM HEPES pH 7.9, 150 mM KCl, 5 mM EDTA, 1 μM maltose) were incubated in 5 mM N-hydroxysulfosuccinimide (Pierce Technology) at room temperature for one hour. Reactions were quenched with 1/10th volume 1 M Tris pH 7.9, and then subjected to SDSPAGE. After staining with Coomassie-G (5% w/v aluminum sulfate 14-18 hydrate, 10% v/v ethanol, 0.02% w/v CBB G-250, and 2% v/v phosphoric acid), the entire protein lane was excised into 6-9 gel slices. For the second acetylation step, gel slices were incubated with shaking agitation under the following sequence of conditions: 1) water for 10 minutes at 37°C, 2) 10 mM ammonium bicarbonate at room temperature, 3) 50 mM ammonium bicarbonate:acetonitrile (1:1) for 45 minutes at 37°C twice, 4) water twice, and 5) 100% acetonitrile at 37°C for 5 minutes twice. After complete removal of the acetonitrile, 20 μl of D6-acetic anhydride (Acros Organics) mixed with 40 μl of 100 mM ammonium bicarbonate was added and completely absorbed by the gel. The gel slices were then submerged in 100 mM ammonium bicarbonate, and the pH adjusted to 7-8 with 1 M ammonium bicarbonate. After 60 minutes at 37°C, the gel slices were washed with water three times and submitted for LCMS/MS analysis.

### Protein digestion and mass spectrometric analysis

For protein digestion, gel pieces were washed in 25 mM ammonium bicarbonate / 60% v/v acetonitrile. Disulfide bonds were reduced using 10 mM dithiothreitol in 25 mM ammonium bicarbonate for 30 minutes at 60°C, and then free sulfhydryls were alkylated using 20 mM iodoacetamide in 25 mM ammonium bicarbonate for an hour. Gel pieces were washed again using 25 mM ammonium bicarbonate / 60% v/v acetonitrile, and then digested overnight using 200 ng TPCK-treated porcine trypsin (Promega) in 25 mM ammonium bicarbonate. Peptides were extracted using 50% v/v acetonitrile/ 1% v/v formic acid. Extracted peptides were dried down by vacuum centrifugation, and resuspended in 0.1% v/v formic acid prior to mass spectrometric analysis.

Peptides were analyzed by LC-MS/MS using a NanoAcquity (Waters) UPLC system interfaced with either an LTQ Orbitrap Velos or QExactive Plus (both Thermo) mass spectrometer. Peptides were separated using an integrated 75 μm x 15 cm PEPMAP reverse phase column and spray tip (EasySpray source). For *E. coli* ribosome samples, a gradient from 2 to 30% v/v solvent B (0.1% v/v formic acid in acetonitrile) over solvent A (0.1 % acetonitrile in water) at a flow rate of 400 nL/min for 72 min was used, followed by ramping to 50% B over 2 min, before returning to starting conditions. For spliceosome samples, a gradient from 2 to 27% v/v solvent B (0.1% v/v formic acid in acetonitrile) over solvent A (0.1 % acetonitrile in water) at a flow rate of 400 nL/min for 27 min was used.

With the Velos, survey scans were measured at a resolution of 30,000 full width at half maximum (FWHM). The six most intense precursor ions were automatically selected for HCD fragmentation analysis by data-dependent acquisition and measured at a resolution of 7,500 FWHM. For QExactive data, survey scans were measured at a resolution of 70,000 FWHM, and were followed by data-dependent acquisition of the top ten most intense precursors at a resolution of 17,500 FWHM.

Raw data were converted to mgf format peak list files using in-house software based on the Raw_Extract script in Xcalibur version 2.4.For ribosome samples, the data were searched with Protein Prospector version 5.21.23 against a database of all *E. coli* proteins in SwissProt downloaded on May 18^th^ 2017, plus decoy versions of all of these sequences (a total of 23,044 entries searched). For spliceosome samples, the data were searched using Protein Prospector version 5.18.22 against a database of all human proteins in SwissProt downloaded on May 9^th^ 2016 supplemented with entries for *E. coli* maltose-binding periplasmic protein, enterobacteria phage MS2 coat protein and pig trypsin, plus decoy versions of all of these sequences (a total of 20,204 entries searched). Searches were performed with tolerances of +/-15 ppm on precursor ions and +/-30 ppm on fragment ions. Carbamidomethylation of cysteine residues was searched as a constant modification, and variable modifications considered were acetylation or deuteroacetylation of uncleaved lysines, methionine oxidation, pyroglutamate formation from N-terminal glutamines, and protein N-terminal methionine removal and/or acetylation. Results were threshholded to an estimated 1% protein FDR according to target:decoy database searching (29). Intensities of peaks were extracted from the raw data using Protein Prospector by summing together signal over a period −10 to +20 seconds from the time the peak was selected for MS/MS.

### Assignment of lysine modifications

Mass spectrometry data for each sample were returned as a list of sequenced peptides with associated protein identifiers.The lists were first filtered for peptides of proteins characterized as stable components of the associated RNP complex (Supplemental Table 1), and then for peptides containing modified lysine residues. From these peptides a catalog of lysine residues was compiled, which were flagged as acetylated (A) or deutero-actylated (D), along with the number peptides supporting the modification state (Supplemental Table 2). In cases where both modifications were identified in an individual sample, we compared the mass spectrometry peak intensity for the parent peptides. In cases where less than 5-fold enrichment for a particular modification was observed, the lysine was flagged as mixed (M).

We next compared results for replicate experiments and initially flagged lysines as A or D in cases where all measurements were in agreement, and M if opposing modifications were present. To estimate the likelihood of mixed modifications arising from inefficiency in probing vs. multiple lysine conformations, we calculated conditional probabilities for three lysine conformations (solvent accessible, buried, and equally present in both states) using Bayes theorem formulated as:

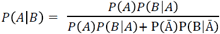

Prior probability P(A) was designated as the frequency of peptides identified with A or D modification, and assuming a low probability (1%) of mixed conformations.Posterior probability P(B|A) for each conformation was calculated as binomial probability mass function for number of specific modification observations per total observations across replicas with modification likelihood based on an estimated probing efficiency of 94% for both steps. This efficiency was derived from the relative frequency of peptides ending in unmodified lysine vs. arginine in the MS data. P(Ā) is 1-P(A), and P(B|Ā) is the sum of posterior probabilities of the other two conformations.These values are reported in Supplemental Table 2.

### Analysis of lysine modification relative to ribosome structure

Structural analyses and generation of molecular graphics were performed with UCSF Chimera (30) and ChimeraX (31), which are developed by the Resource for Biocomputing, Visualization, and Informatics at the University of California, San Francisco, with support from NIH R01-GM129325 and P41-GM103311. Lysine modification data for *E. coli* ribosomes was compared to the structure of the 30S subunit extracted from RSCB entry 4YBB. Distances from lysine primary amines to their nearest neighbor were determined with the “findclash” command. Solvent accessible surface area for the primary amine was calculated with the FreeSASA program (32). B-factors were extracted from the PDB file. The values reported are average measurements from the two 30S subunits in the crystal’s asymmetric unit. Data were plotted using ggplot2 in RStudio.

## RESULTS

### Two-step probing methodology and lysine acetylation assignment

Lysine side chains are good targets for chemical probing because they are reactive to acetylation reagents under mild conditions that maintain molecular interactions. The modification state of an individual lysine residue can then be determined by LC-MS/MS of peptides derived from proteins in the macromolecular complex that was subjected to chemical probing. However, a caveat to such analysis is that peptides for mass spectrometry evaluation are routinely generated by tryptic digestion, which cleaves after unmodified lysine and arginine residues, but not after acetyl-lysine. Assessing changes in lysine modification states between multiple RNP conformations is problematic because differential acetylation patterns will generate different tryptic peptides that are not directly comparable by MS/MS due to varying retention times and ionization profiles. To overcome this challenge, we developed a two-stage lysine acetylation protocol in which the native RNP is first probed with sulfo-NHS, and then denatured by SDS-PAGE before probing a second time with deuterated acetic anhydride (Figure 1A-B). Because every lysine present in the protein should be modified in either the first or second step, the strategy will yield identical tryptic peptides. The 3 mass-unit difference between acetyl-lysine and deutero-acetyl-lysine allows us to discriminate between reactivity in the RNP vs. the denatured state (Figure 1C).

**Figure 1:**
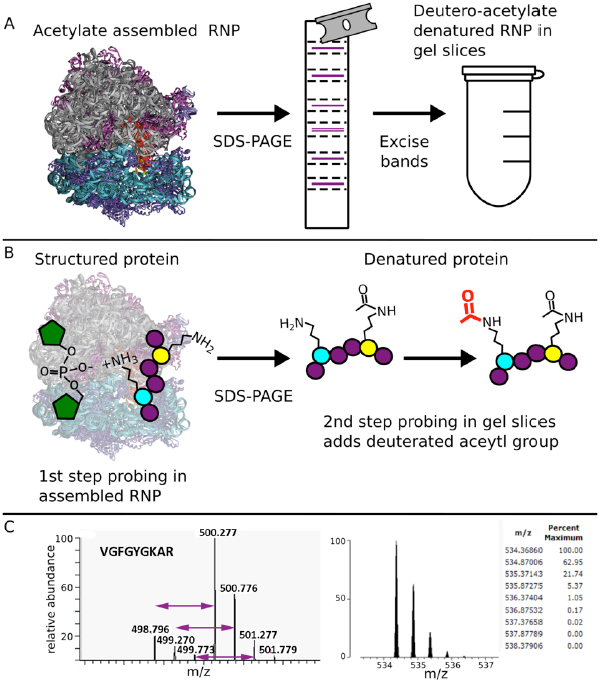
Two-step acetylation protocol. A) Cartoon diagram of an assembled RNP (70S ribosome) being acetylated, denatured, the deutero-acetylated. B) Cartoon diagram showing a lysine interacting with the negative backbone of RNA and thus is resistant to the first acetylation step being deutero-acetylated (red acetyl group) and a lysine residue sensitive to the first acetylation (black acetyl group). C. Left: Sample MS spectra containing two doubly charged peptides of m/z 498.796 and m/z 500.277 corresponding to the peptide VGFGYKAR from *E. coli* protein RS5. In the m/z 498.796 version, the lysine is modified with an acetyl group, whereas in the m/z 500.277 version, the lysine is modified with a deuteron-acetyl group, leading to a 3 Dalton mass difference. Right: Theoretical modeling of the predicted isotope pattern for the lighter version using MS-Isotope (34) indicates a fourth isotope peak would contribute 5% peak intensity to the first isotope of the deuterated monoisotopic peak signal, and have an insignificant effect on overall quantification

We tested the method by probing two different RNPs: the *E. coli* 30S ribosome subunit and human spliceosomes. Because of the limited material obtainable for many macromolecular assemblies, such as *in vitro* assembled spliceosomes, we analyzed low picomole amounts of complexes for both sample types. After two-step chemical probing, MS/MS analysis of individual samples generated lists of sequenced peptides and modifications (Supplemental Table 1). Across ten samples analyzed (three replicates each of intact 30S ribosome subunit and of isolated 30S proteins, and duplicate preparations of two spliceosome conformations), >96% of lysine residues present in the MS/MS peptide sequences were reported as modified (Table 1). Correspondingly, >96% of peptides obtained from trypsin digest ended in arginine or the final amino acid of the protein. Assuming equal modification rates at each step and considering that any accessible residue in the RNP that escaped modification in the first step would likely be modified in the second step, we estimate probing efficiency for each reaction to be above 94%. Importantly, the high rate of modification produced a large number of comparable peptides between replicates and between different conformations of the same complex.

**Table 1:**

| Sample type | Replicate | MS/MS peptides |  | Modified lysines |  |  | Modification state |  |  |
| --- | --- | --- | --- | --- | --- | --- | --- | --- | --- |
|  |  | From complex | Ending in K | Observed | Measured >1 time | With same modification | %A | %D | %M |
| 30S ribosome subunit | 1 | 136 | 0% | 100 | 31 | 74.2% | 30.0 | 62.0 | 8.0 |
|  | 2 | 196 | 8.1% | 135 | 61 | 86.9% | 34.1 | 58.5 | 7.4 |
|  | 3 | 151 | 3.6% | 111 | 46 | 82.6% | 31.5 | 61.3 | 7.2 |
| 30S ribosome proteins only | 1 | 190 | 4.5% | 111 | 43 | 74.4% | 67.6 | 22.5 | 9.9 |
|  | 2 | 199 | 4.9% | 110 | 50 | 60.0% | 57.3 | 24.5 | 18.2 |
|  | 3 | 57 | 8.9% | 35 | 11 | 81.8% | 77.1 | 17.1 | 5.7 |
| Catalytic spliceosome | 1 | 847 | 0% | 489 | 126 | 79.4% | 66.5 | 29.2 | 4.3 |
|  | 2 | 691 | 1.8% | 319 | 81 | 69.1% | 33.5 | 57.1 | 9.4 |
| Post-catalytic spliceosome | 1 | 549 | 0% | 329 | 104 | 44.2% | 61.7 | 22.2 | 16.1 |
|  | 2 | 1068 | 1.1% | 466 | 181 | 87.8% | 57.7 | 26.6 | 15.7 |
| Average |  |  | 3.3% |  |  | 74.7% |  |  |  |

For each sample, we next compiled a list of individual lysine residues, which we flagged as acetylated (A) or deutero-acetylated (D) based on the associated modification. We found that within an individual sample, certain peptides were reported multiple times when they carried additional modifications, such as oxidized methionine. When also containing a lysine, these peptides served as independent measurements of the modification state, and nearly 70% carried the same modification. In the cases for which a given lysine was identified in multiple peptides with both acetyl and deutero-acetyl modification, we compared the mass spectrometry peak intensity for the parent peptides. If the ratio indicated enrichment of less than five-fold for a particular modification, we flagged the lysine as mixed (M). Mixed modifications may be due to incomplete acetylation of the native RNP, but a comparison of modifications between replicates, as discussed below, suggests that more often they reflect heterogeneity in the composition or conformation of the RNP being probed.

We assessed probing reproducibility by comparing three replicate experiments for the 30S ribosome subunit. As a result of the two-step strategy, over half of the MS peptides measured overall were present in all three replicates, and 118 individual lysine residues were observed more than once (Figure 2A). Of these, we found that 78% exhibited the same modification state across replicates (Figure 2B). The remaining were identified as mixed in one or more samples, or exhibited opposing modifications (A vs. D). Notably, mixed and opposing modifications were not randomly distributed among all residues, as would be expected if inefficient probing alone were the cause.

**Figure 2:**
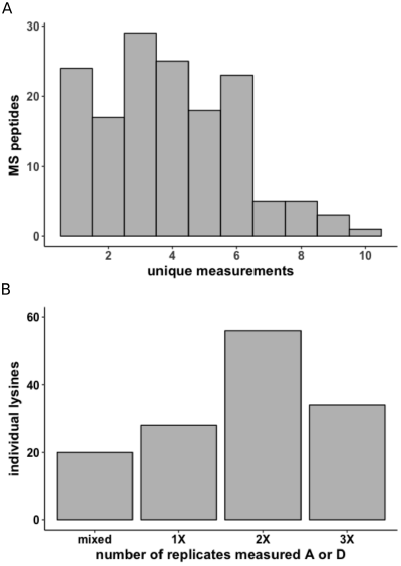
30S ribosome subunit two-step acetylation reproducibility A) Frequency at which MS peptides are uniquely measured B) MS lysine modification reproducibility across replicates.

To better understand the source of mixed modifications, we calculated conditional probabilities for three possible lysine orientations in the 30S ribosome subunit: solvent exposed, solvent inaccessible, and equally present in both conformations. Using the estimated modification efficiency of 94% at both probing steps, we compared a model in which probing inefficiency accounted for nearly all our observations of mixed modification to a model in which multiple conformations of the lysine was the primary cause. The analysis indicated that 80% of differentially modified lysines were more likely due to mixed conformations present in the sample, rather than to probing inefficiency. We used these probabilities to ultimately classify lysines as primarily solvent accessible (A), primarily buried (D) or present in both conformations (M) (Supplemental Table 2).

### Comparing lysine probing patterns of the 30S ribosome subunit to crystallographic structures

We mapped our results on a high-resolution model of the 30S *E. coli* ribosome subunit derived from X-ray crystallography (33) to assess the consistency of lysine probing patterns with structural models of RNPs. In the structure, a majority of deutero-acetyl-lysines are modeled in conformations consistent with ionic interactions with the phosphate backbone of rRNA, while lysines with acetyl or mixed modification state are generally directed toward solvent (Figure 3). To more quantitatively evaluate the relationship between lysine reactivity and structure, we also compared the observed modification state to three parameters: solvent accessible surface area (SASA), distance to the nearest atom and B-factor, all relative to the primary lysine amine derived from the same ribosome crystal structure (Figure 4A-C, Supplemental Table 3). Consistent with the primary amine being constrained by interactions that reduce lysine reactivity in the native ribosome, values for deutero-acetyl-lysine trended lower for these three parameters relative to acetyl-lysine and mixed.

**Figure 3:**
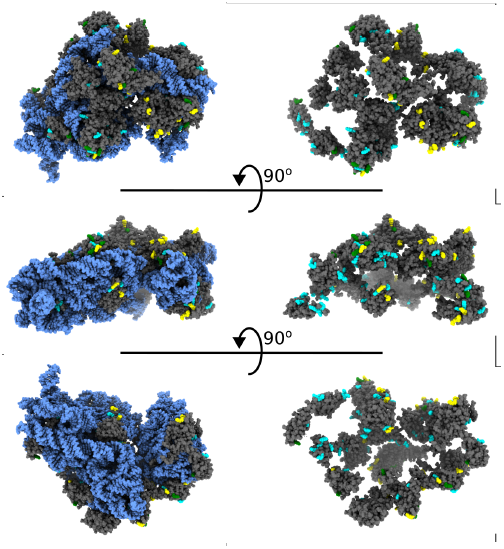
MS/MS lysines colored to match their acetylation state displayed on the 30S subunit of the 70S ribosome. Blue is 16S RNA, grey are 30S proteins, cyan is deutero-acetyl lysines, yellow is acetyl lysine and green are mixed.

**Figure 4:**
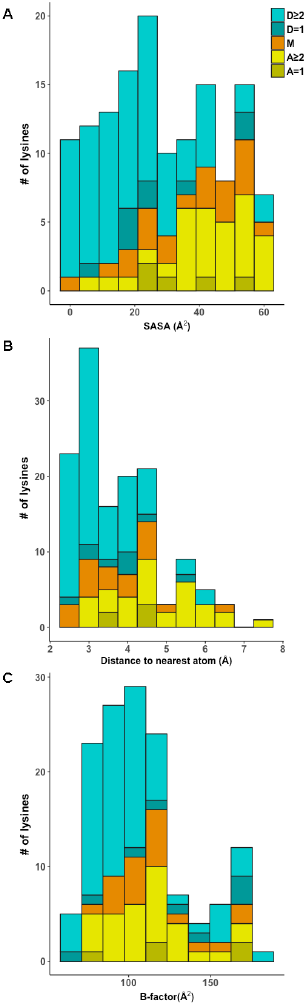
Stacked histograms of A) SASA scores, B) distance to nearest atom and C) B-factor of individual MS/MS lysines separated by modification state and the number of times a given lysine was measured across replicates. A is acetyl-lysine, D is deutero-acetyl-lysine and M is mixed. Shading indicates whether the lysine was observed in one (=1, darker) or more than one (≥2, lighter) replicates.

To further test the relationship between two-step probing results and molecular structure, we modified isolated 30S ribosome proteins. As expected, in the absence of the 16S rRNA, lysines from these proteins were identified with a higher proportion of acetyl modifications relative to the intact RNP (Figure 5). We also saw an increased proportion of lysines likely in mixed conformations, which may be due to conformational heterogeneity of the predicted intrinsically disordered regions in most ribosome proteins in the absence of rRNA. Still, 80% of lysines in free 30S proteins exhibit the same modification state across replicates (Supplemental Table 4). Together, these data indicate our probing strategy can discern the loss of 30S protein interactions in the absence of 16S rRNA, and indicates that the proteins largely maintain a consistent structure in terms of lysine accessibility.

**Figure 5:**
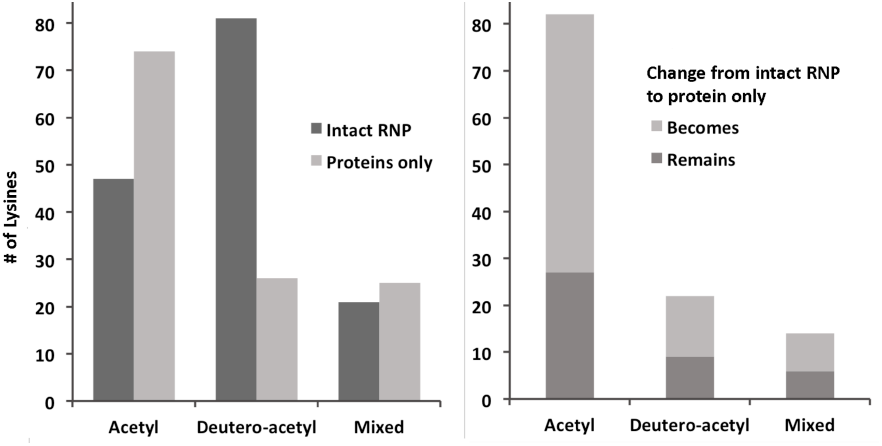
Modification changes between intact 30S ribosome subunit versus isolated 30S ribosome proteins only.Left panel:Number of lysines with indicated modification states.Right panel: Number of lysines that retain or change to the indicated modification state in isolated 30S ribosome proteins relative to the intact RNP.

### Comparing multiple conformations of the spliceosome

In addition to indicating the local structural environment of a lysine residue, chemical probing should indicate similarities and differences in that environment between different functional conformations of a macromolecular complex. We probed different forms of human spliceosomes isolated from *in vitro* splicing reactions, and again many identical peptides resulted from the two-step protocol (Supplemental Table 1). Comparing catalytic and post-catalytic human spliceosomes, 69% of the 404 lysines observed in both samples had the same modification (Supplemental Table 4). Some proteins shared between the complexes showed no differences at all, while others had more modification changes (Figure 6A). This result is consistent with cryo-EM models of similar states of yeast spliceosomes, which exhibited a high degree of similarity in overall architecture (3). Lysines that change in accessibility as the spliceosome carries out exon ligation may indicate interaction sites for incoming / outgoing components or reflect conformational changes that regulate spliceosome progression.

**Figure 6:**
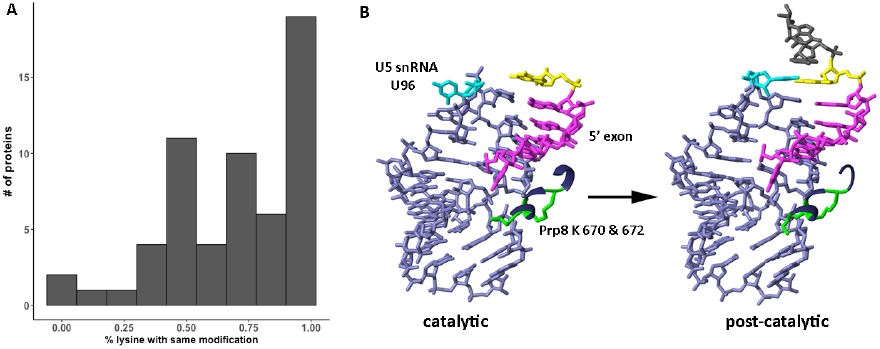
Modification changes between catalytic and post-catalytic spliceosomes.A.Histogram depicting the distribution of the percentage of overall modification differences for the 58 proteins present in both complexes. B. Close-up view of the relative positions of U5 snRNA stem-loop I (lavender), 5 exon (magenta), and differentially modified Prp8 lysine residues K670 and K672 (green) in catalytic and post-catalytic spliceosome conformations. The last nucleotide of the 5’ exon is highlighted in yellow, and U96 of U5 snRNA is in cyan.

We recently reported evidence for a transient conformational change in the spliceosome before and after activation based on two-step chemical probing (26). A change in modification state for two conserved lysine residues in the core scaffold protein Prp8 indicated that they gain contacts during formation of the active site. In cryo-EM models of both states, these lysines are modeled in close proximity to the U5 small nuclear RNA (snRNA) near its interaction with the 5’ exon of the pre-mRNA substrate (2,4,5,16,21). Guided by the probing results, we closely examined the relative positions of U5 snRNA and the 5’ exon between preactivated and catalytic spliceosomes, and identified a change in register that suggests transient conformation involving changes in the lysine interactions. Importantly, genetic experiments in yeast guided by the probing data confirmed a functional role for the lysines in correctly positioning the 5 exon of the pre-mRNA for catalysis. Notably, we now see that the same two lysines (Prp8 670K and 672K) become more reactive with the transition to the post-catalytic state, further suggesting an inherent flexibility in this region. Although these lysines are modeled in essentially the same location in the cryo-EM models of the catalytic and post-catalytic spliceosomes, their change in reactivity may be linked to a movement in nearby U96 of U5 snRNA, which flips toward the active site in the post-catalytic complex to interact with the last nucleotide of the exon (Figure 6B)(15,16). Together our data illustrates how chemical probing can contribute to identifying and characterizing transient interactions that may not be easily distinguished in static structural models of macromolecular assemblies.

## DISCUSSION

Technological advances in cryo-EM have brought structure determination of many multi-megadalton biological assemblies finally within reach. Now, more than ever, interpretation of these models needs testing to support or reject putative molecular mechanisms suggested by structural architecture. This need is further intensified by the knowledge that macromolecular complexes in aqueous solution often sample multiple conformations.Many of these conformations will not be represented in molecular models generated from cryo-EM or X-ray crystallography, which are derived from averaging over molecules imbedded in vitreous ice or situated in a crystalline lattice, respectively. While a cryo-EM preparation theoretically includes all conformations of the molecule that exist at the time of rapid freezing, non-abundant conformations are often lost as a result of image processing. In contrast, chemical probing reports on all aqueous conformations, and can add an additional layer of information for characterizing flexible regions of proteins and nucleic acids, which are relevant to their function.

The two-step chemical probing method described here specifically measures changes in lysine interactions. The method is useful when assessing structural variations of proteins in solution, even in the context of dynamic entities like the spliceosome. Given the continuous negative charge of a RNA backbone, lysine probing is particularly helpful in analyzing RNPs, but can also be applied to any molecular complex. When employing the methodology, a few points should be considered. First, low-picomole amounts of material are sufficient to capture a large portion of the sequence of a given protein, but more material will allow for greater coverage, especially if used in conjunction with other proteases for generating peptides for the MS/MS analysis. Second, for all of the data presented here, the first probing step of native complexes occurred over the course of an hour. Because of this, interactions that quickly fluctuate between two states are likely to be identified as solvent accessible. A potential extension of two-step probing would be to carry out a time course during the first probing period to differentiate more dynamic lysine residues. Third, more sophisticated mass spectrometry strategies, like multiple reaction monitoring, could be used to better quantify the relative ratios of differentially modified lysines. Finally, modifying reagents against other amino acids can be additionally incorporated into the protocol if their respective chemistries occur at a pH at which the target complex is purified.

## SUPPLEMENTARY DATA

Supplementary Data are available at NAR online.

## ACKNOWLEDGEMENT

We thank Laura Lancaster and Harry Noller for providing reconstituted 30S *E. coli* ribosome subunits and isolated 30S proteins.

### FUNDING

This work was funded by National Institutes of Health (R01GM72649 to M.S.J.; RR001614, RR015804, RR019934, P41 GM103481 to R.J.C. and A.L.B.) and the Adelson Medical Research Foundation (to A.L.B). A.J.M. was supported by National Institutes of Health Training Grant T32GM08646.P.C. was supported by FAPESP grant 2017/06994-9 and CNPq grant 474672/2013-1.

### CONFLICT OF INTEREST

The authors declare no conflicts of interest.

